# Near-cognate initiation generates FMRpolyG from CGG repeats in Fragile X associated Tremor Ataxia Syndrome

**DOI:** 10.1101/2020.10.19.345736

**Authors:** Yuan Zhang, M. Rebecca Glineburg, Venkatesha Basrur, Kevin Conlon, Deborah A. Hall, Peter K. Todd

## Abstract

Repeat associated non-AUG (RAN) translation of *FMR1* 5’ UTR CGG repeats produces toxic homo-polymeric proteins that accumulate within ubiquitinated inclusions in Fragile X-associated tremor/ataxia syndrome (FXTAS) patient brains and model systems. The most abundant RAN product, FMRpolyG, initiates predominantly at an ACG codon located just 5’ to the repeat. Methods to accurately measure FMRpolyG in FXTAS patients are lacking. Here we used data dependent acquisition (DDA) and parallel reaction monitoring (PRM) mass spectrometry coupled with stable isotope labeled standard peptides (SIS) to identify potential signature FMRpolyG fragments in patient cells and tissues. Following immunoprecipitation (IP) enrichment, we detected FMRpolyG signature peptides by PRM in transfected cells, FXTAS human samples and patient derived stem cells, but not in controls. Surprisingly, we identified two amino-terminal peptides: one beginning with methionine (Ac-MEAPLPGGVR) initiating at an ACG, and a second beginning with threonine (Ac-TEAPLPGGVR), initiating at a GUG. Abundance of the threonine peptide was enhanced relative to the methionine peptide upon activation of the integrated stress response. In addition, loss of the eIF2 alternative factor, eIF2A, or enhanced expression of initiation factor eIF1, preferentially suppressed GUG initiated FMRpolyG synthesis. These data demonstrate that FMRpolyG is quantifiable in human samples and that RAN translation on *FMR1* initiates at specific near cognate codons dependent on available initiation factors and cellular environment.

## Introduction

Fragile X associated tremor ataxia syndrome (FXTAS) is an inherited late-onset neurodegenerative disorder caused by a CGG repeat expansion within the 5’UTR of *FMR1*. One of the pathological hallmarks of this disease is the presence of ubiquitinated neuronal inclusions in patient brain tissue (Greco et al. 2002; Greco et al. 2006). These inclusions are thought to occur as a result of an aberrant form of translation, termed repeat associated non-AUG (RAN) translation (Zu et al. 2011; Todd et al. 2013; Glineburg et al. 2018). RAN translation on *FMR1* transcripts is enhanced by CGG repeat expansions that form strong secondary structures in the RNA (Kearse et al. 2016; Krzyzosiak et al. 2012; Zumwalt et al. 2007; Verma et al. 2020). These repeats are thought to stall scanning ribosomes, allowing initiation just upstream of or within the repeat in the absence of an AUG start codon (Kearse et al, 2016). RAN translation at CGG repeats (CGG RAN) can occur in all three reading frames, producing three distinct toxic homopolymeric peptides, the most abundant of which is a polyglycine product, FMRpolyG (Todd et al. 2013; Kearse et al. 2016; Green et al. 2017; Linsalata et al. 2019). Overexpression of CGG repeats that support FMRpolyG expression promotes neurodegeneration in various model systems (Todd et al. 2013; Sellier et al. 2017; Oh et al. 2015; Rodriguez et al. 2020; Linsalata et al. 2019; Hukema et al. 2015; Hoem et al. 2019) and FMRpolyG is detected in human FXTAS brain tissue where it localizes to ubiquitin and P62+ inclusions (Krans et al. 2019; Buijsen et al. 2016; Buijsen et al. 2014). FMRpolyG has recently been detected via mass spectrometry (MS) in transfected cells as well as in patient tissue (Sellier et al. 2017; Ma et al. 2019); however, reliable detection of CGG RAN peptides and the ability to accurately quantify them is lacking.

In reporter assays, RAN translation of FMRpolyG predominantly initiates at two near cognate codons: an ACG, located 36 nts upstream of the repeat and a GUG located 12 nts from the repeat (Kearse et al. 2016; Sellier et al. 2017). Exactly how this initiation occurs and what factors are required is not fully understood. Recently, our lab and others uncovered a role for the integrated stress response (ISR) in upregulating RAN translation at CGG repeats and at the *C9orf72* ALS-causing G_4_C_2_ repeat (Green et al. 2017; Cheng et al. 2018; Sonobe et al. 2018; Westergard et al. 2019). Under normal conditions, eIF2 binds GTP and Met-tRNA_i_^Met^ to form the ternary complex (TC) that then binds the 40S ribosome and a host of other translation initiation factors that facilitate cap binding and scanning, to form the 43S preinitiation complex (PIC). The PIC then scans along the mRNA, 5’-3’, until it encounters the AUG start codon and initiation occurs (Sonenberg and Hinnebusch 2009; Hinnebusch 2011; Jackson, Hellen, and Pestova 2010; Hershey 2000). However, when any one of four ISR kinases (PERK, GCN2, HRI, and PKR) are activated, they phosphorylate canonical translation initiation factor eIF2α, thereby inhibiting eIF2β from exchanging GDP for GTP, and preventing a new round of TC assembly. This effectively inhibits canonical translation as no new TCs can be loaded onto 40S ribosomes to promote initiation (Jackson, Hellen, and Pestova 2010; Walter and Ron 2011).

Despite this global inhibition, some genes are able to bypass this response, and many do so by initiating at near-cognate codons, rather than the canonical AUG start codon (Starck et al. 2016; Ingolia et al. 2009; Green et al. 2017; Imataka, Olsen, and Sonenberg 1997; Ingolia, Lareau, and Weissman 2011; Kearse et al. 2019; Schwab et al. 2004). The likelihood that initiation will occur at a near-cognate codon during cellular stress is dependent on a number of factors including efficiency of TC recruitment to the 40S docked on an initiation codon, strength of that codon and its surrounding Kozak sequence, and availability of specific initiation factors (Kolitz, Takacs, and Lorsch 2009; Lind and Aqvist 2016; Tang et al. 2017; Kearse and Wilusz 2017). In particular, the eIF2 alternative factor, eIF2A, can facilitate near-cognate codon initiation, and a unlike eIF2, is GTP-independent and upregulated during cellular stress (Starck et al. 2016; Starck et al. 2012; Komar et al. 2005; Kim et al. 2018; Sonobe et al. 2018). eIF2A has also been implicated in RAN translation of G_4_C_2_ repeats that initiates at a CUG codon (Sonobe et al. 2018).

While eIF2A is capable of promoting non-AUG initiation, the canonical initiation factor eIF1 has an opposing role of ensuring AUG start-codon fidelity. eIF1 does this by 1) keeping the 43S PIC in an opened conformation to allow for TC binding and scanning, 2) impeding eIF5 GTPase activity at non-AUGs, which slows down the conversion of eIF2-GTP to eIF2-GDP-Pi, and inhibits Pi release, and 3) dissociating from the ribosome only after the PIC has bound to an AUG (Maag et al. 2005; Algire, Maag, and Lorsch 2005; Passmore et al. 2007; Unbehaun et al. 2004). In vitro experiments have shown that in the absence of eIF1, ribosomes can readily initiate at near cognate codons, and when eIF1 is overexpressed, use of near cognate codons is severely inhibited, specifically when the near-cognate codon differs at position 1 (Valasek et al. 2004; Pestova and Kolupaeva 2002; Fekete et al. 2007; Lind and Aqvist 2016). eIF1 overexpression was recently shown to dramatically inhibit CGG RAN translation (Linsalata et al. 2019). How eI2A and eIF1 might interact to dictate start codon selection and whether or how they contribute to CGG RAN remain unknown.

Here, using parallel reaction monitoring (PRM) MS coupled with stable isotope labeled standard peptides (SIS), we both detected and quantified FMRpolyG peptides in overexpression systems and in FXTAS patient cells and samples. Furthermore, we identified an additional initiation codon, GUG, immediately 5’ of the ACG, that is preferentially used in response to cellular stress. Initiation at this codon was selectively inhibited in the absence of eIF2 alternative factor, eIF2A, and during overexpression of canonical translation initiation factor, eIF1. Together these data provide a promising new technique for quantifying low abundant RAN peptides, and provide mechanistic insights into non-AUG initiation in a neurodegenerative disease.

## Results

### Development of PRM-SIS for FMRpolyG quantification

RAN translation of FMRpolyG primarily initiates at two near cognate codons, ACG and GUG, located in the *FMR1* 5’ UTR, upstream of the CGG repeat (Kearse et al. 2016); however, while initiation at the ACG with a methionine has been previously reported in transfected samples, and recently found at extremely low levels endogenously, whether other species exist, and whether they can be detected at quantifiable levels has not been determined (Sellier et al. 2017; Ma et al. 2019). In order to detect and quantify endogenous FMRpolyG, we first identified potential FMRpolyG signature peptides. We predicted four different FMRpolyG sequences based on known initiation start sites for FMRpolyG: two longer isoforms beginning at ACG and initiating with either a methionine or a threonine, and two shorter isoforms beginning at GUG and initiating with either a methionine or a valine (Figure 1A). From these sequences, we identified four potential FMRpolyG signature peptides (two N-terminal and two C-terminal) to use for quantification (Figure 1A underlined sequences). We then applied data dependent acquisition (DDA) MS to identify these signature peptides in an FMRpolyG overexpression system. After comparing with the latest SwissProt human protein database (42054 sequences) (https://www.uniprot.org/statistics/Swiss-Prot), we found three of the four signature FMRpolyG peptides produced in HEK293Ts expressing exogenous FMRpolyG: the two C-terminal fragments——a carbamidomethyl modified peptide, CAM-CGAPMALSTR, and an unmodified peptide, SPPLGGGLPALAGLK——and an acetyl-modified methionine-initiated N-terminal fragment, Ac-MEAPLPGGVR (Figure 1B-D). This N-terminal fragment and the SPPLGGGLPALAGLK fragment have previously been identified using similar methods (Sellier et al. 2017; Ma et al. 2019).

**Figure 1.**
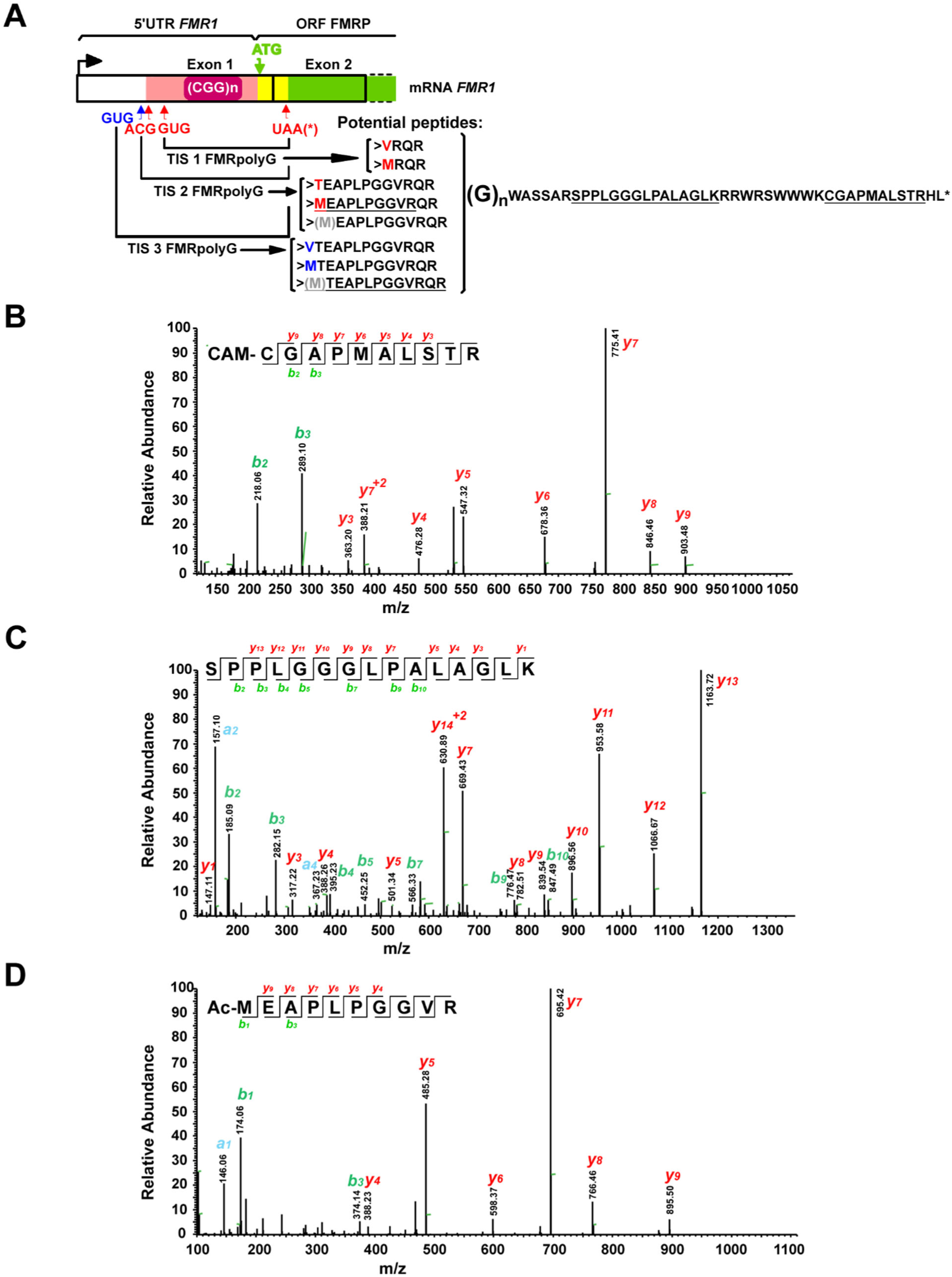
Predicted potential peptide sequences for RAN translation product FMRpolyG. **A**. Schematic of the peptides produced via alternative translation initiation on *FMR1*. Top) *FMR1* schematic showing location of the FMRpolyG uORF (pink), FMRP ORF (green), and sequence corresponding to overlap of these two ORFs (yellow). Bottom) The predicted FMRpolyG peptide sequences from three potential translation initiation sites (TIS) on *FMR1*, two previously identified initiation sites (red), and one novel initiation site (blue). Predicted trypsin-digested signature peptides for FMRpolyG that are monitored in this study are underlined. The grey M in brackets indicates a potential product of iMet excision. **B-D**. LC-MS/MS spectra of FMRpolyG signature peptides CAM-CGAPMALSTRHL (B), SPPLGGGLPALAGLK (C) and Ac-MEAPLPGGVR (D) from HEK293T cells transfected with FMRpolyG_100_-3XFlag. Observed *b-* and *y-ions* are indicated.

The carbamidomethyl modification on an NH_2_-terminal cysteine can cause the peptide to spontaneously undergo cyclization during MS sample preparation in a pH- and time-dependent manner, making CAM-CGAPMALSTR a poor peptide for quantification (Geoghegan et al. 2002). Thus, we chose Ac-MEAPLPGGVR and SPPLGGGLPALAGLK as signature peptides for FMRpolyG quantification by parallel reaction monitoring combined with stable isotope-labeled signature peptides (PRM-SIS). PRM-SIS is one of the most accurate approaches for directly quantifying the absolute abundance of known target proteins in a complex mixture (Peterson et al. 2012). By monitoring the response and determining the area under the curve of specific fragment ions of both endogenous and SIS peptides (spiked in at known concentrations during sample preparation), we can calculate the absolute abundance of our protein in a given sample (Figure 2A). Light and heavy peptides for Ac-MEAPLPGGVR and SPPLGGGLPALAGLK were synthesized and used to determine the limit of detection and limit of quantification in the background of Fragile X Syndrome patient derived fibroblasts which do not produce any *FMR1* mRNA, as well as in control CSF (Supplementary Figure 1A-D, Supplementary Table 1). Both conditions had values in the sub-nanomolar range (data not shown).

**Figure 2.**
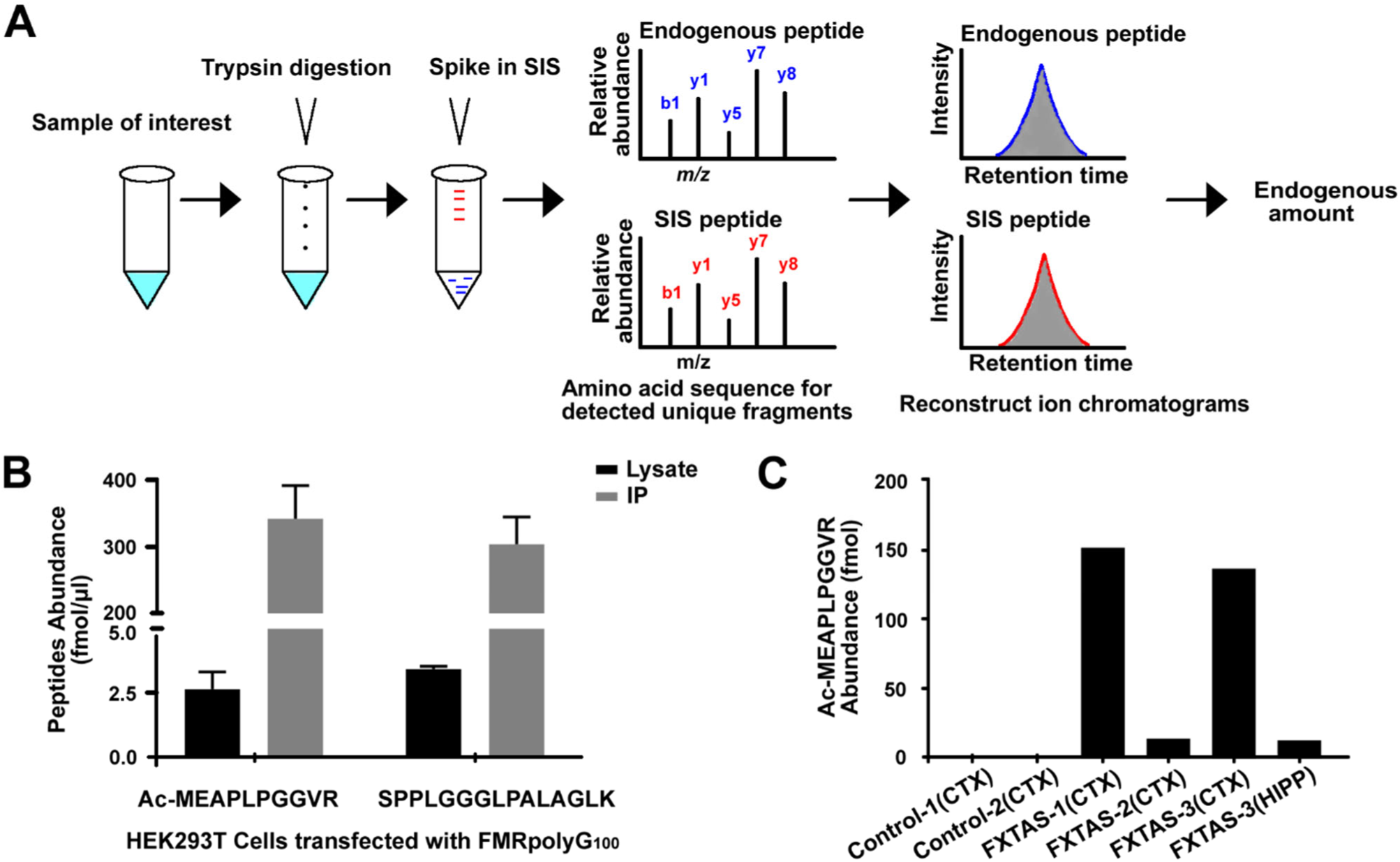
Parallel Reaction Monitoring based quantification of FMRpolyG. **A**. Schematic for PRM-SIS in quantification. Samples were digested with trypsin and then spiked with known concentrations of a signature stable isotope-labeled (SIS) peptide. The amino acid sequence for our signature peptide of interest is detected by MS, and the endogenous peptide concentration is quantified relative to the SIS peptide. **B**. Quantification of signature peptides for FMRpolyG, Ac-MEAPLPGGVR and SPPLGGGLPALAGLK, in HEK293T cell lysates transfected with untagged FMRpolyG_100_ for 24 hrs and either processed directly or following immunoprecipitation with NTF1 antibody. Bars represent mean ± SD. N=2. **C**. Quantification of the absolute abundance of endogenous FMRpolyG peptide Ac-MEAPLPGGVR from human FXTAS and control brains by PRM-SIS.

### Optimizing FMRpolyG detection in transfected HEK293Ts by PRM-SIS

We next quantified FMRpolyG abundance in HEK293Ts transfected with or without untagged FMRpolyG_100_. Despite being able to detect FMRpolyG in overexpression systems quite readily by western blot (Supplemental Figure 2C) (Todd et al. 2013; Kearse et al. 2016; Green et al. 2017), measurement of the two signature peptides Ac-MEAPLPGGVR and SPPLGGGLPALAGLK by PRM-SIS were near the limit of quantification even with overexpression (Figure 2B). This discrepancy was not due to FMRpolyG aggregation, as FMRpolyG was primarily in the soluble fraction following ultracentrifugation, with only a small fraction found in the urea soluble pellet (Supplemental Figure 2A, B). Furthermore, FMRpolyG was efficiently digested by trypsin (Supplemental Figure 2C), suggesting the low quantification observed was not due to insufficient digestion of FMRpolyG but due to low abundance. Consistent with these observations, digesting lysates with the pentaglycine endopeptidase, lysostaphin (LS), which cleaves the polyglycine stretch in FMRpolyG (Todd et al. 2013), had no effect on the levels of Ac-MEAPLPGGVR, and only modestly enhanced detection of SPPLGGGLPALAGLK (Supplemental Figure 2D, E). To overcome this low level of detection, we enriched for FMRpolyG through immunoprecipitation (IP). Following IP of FMRpolyG using a specific FMRpolyG N-terminal antibody (NTF1) (Krans et al. 2019), FMRpolyG was detected >100 fold more than in whole cell lysate (Figure 2B). This fold increase correlated with amounts of starting material (50 μ g/cell lysate vs 4mg/IP) and was maintained when normalized to spiked-in SIS Ac-MEAPLPGGVR and SPPLGGGLPALAGLK.

### Detection and Quantification of endogenous FMRpolyG in FXTAS patient derived samples

We next sought to detect FMRpolyG in IPed human derived samples using PRM-SIS. Only one of the monitored FMRpolyG peptides, Ac-MEAPLPGGVR, was detectable at low levels in a CGG premutation iPSC line (FX 11-9U) and an un-methylated full mutation line (Rodriguez et al. 2020) but not in a control iPSC line with 23 repeats (2E) (Supplemental Figure 4A-B and Supplemental Table 2). We reasoned that the above are not ideal cell types to detect FMRpolyG, since *FMR1* is highly expressed in the brain and most FMRpolyG pathology reported to date is seen in neurons and glia (Tassone et al. 2004; Greco et al. 2006; Iwahashi et al. 2006; Hagerman et al. 2007; Todd et al. 2013). Using PRM-SIS, we detected Ac-MEAPLPGGVR in FXTAS cortex samples from 3 separate cases and 1 FXTAS hippocampus sample, but not in 2 age-matched control cortices (Figure 2C). The absolute abundance for Ac-MEAPLPGGVR ranged widely from 2.96 to 45.36 fmol/mg (Figure 2C, and Supplemental Table 2). We observed more FMRpolyG in cortex vs hippocampus for FXTAS-3, which may reflect the increased diffuse staining (rather than accumulation in ubiquitinated inclusions) observed for cortical FMRpolyG by immunohistochemical techniques (Krans et al. 2019).

### Discovery of an additional RAN initiation codon

DDA MS only detected one N-terminal fragment, Ac-MEAPLPGGVR. However, using the more sensitive PRM method, we identified a second N-terminal peptide—Ac-TEAPLPGGVR (Figure 3A). To measure this peptide, we synthesized light and heavy SIS for Ac-TEAPLPGGVR, and determined the LOD (0.9766 fmol/µl) and LOQ (2.9298 fmol/µl) in Fragile-X Syndrome fibroblasts to be in the sub-nanomolar range similar to that of Ac-MEAPLPGGVR (Supplementary Figure 3A-B). When we quantified Ac-TEAPLPGGVR from HEK293Ts transfected with untagged FMRpolyG, we found the abundance for the new Ac-TEAPLPGGVR fragment of FMRpolyG was lower than the Ac-MEAPLPGGVR fragment (Figure 3B, *see* Figure 2B), but still made up a significant proportion (15.69%) of detected FMRpolyG initiation peptides. We saw a similar pattern in HEK293Ts transfected with FMRpolyG_100_-3xFlag, using lysates directly (17.42%) and following IP (18.37%) (Figure 3D).

**Figure 3.**
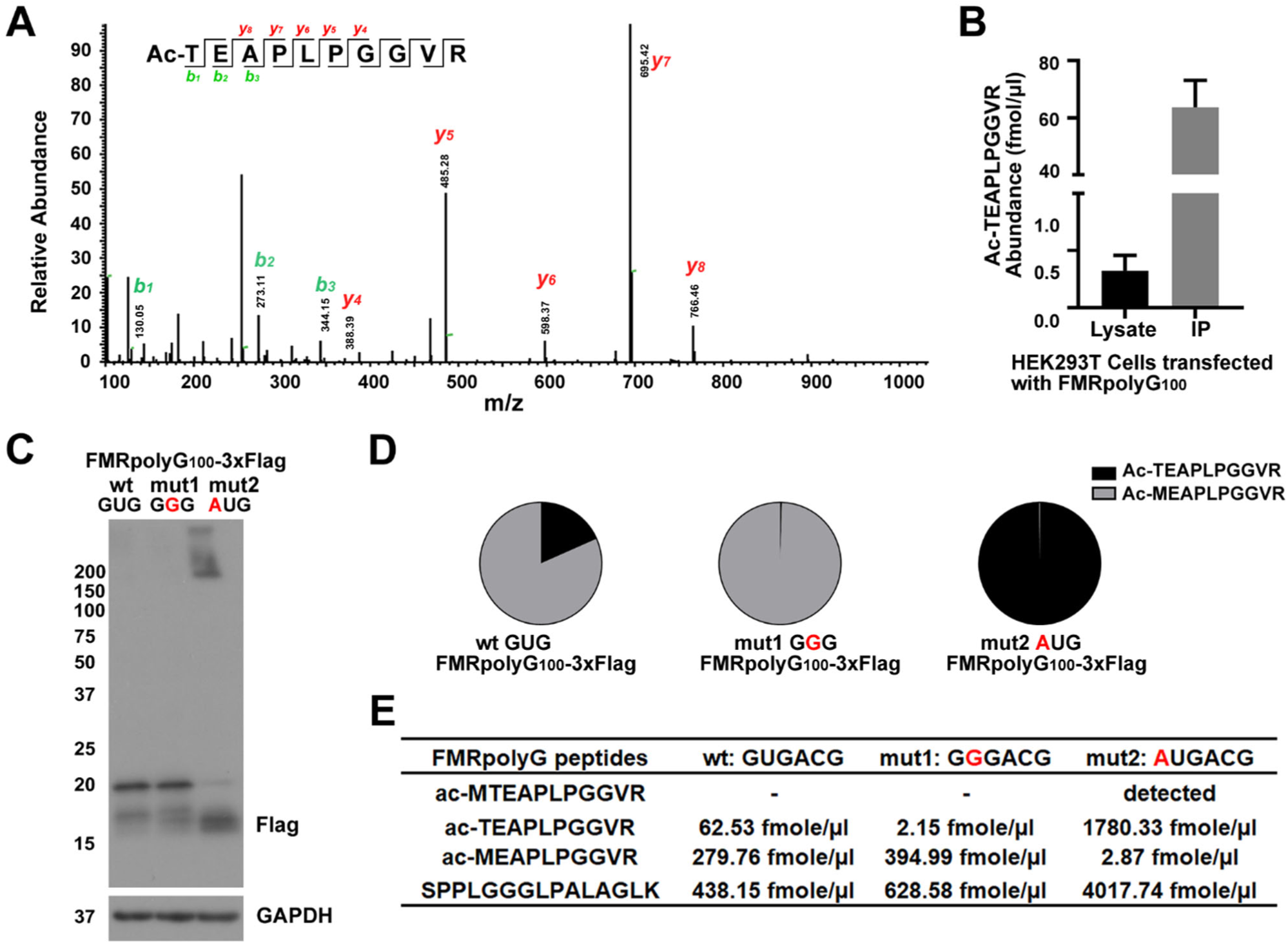
CGG RAN translation initiation can occur at a 5’ GUG codon. **A**. Representative LC-MS/MS chromatogram of FMRpolyG peptide, Ac-TEAPLPGGVR from NTF1 immunoprecipitated HEK293T cells transfected with FMRpolyG_100_. **B**. Quantification of Ac-TEAPLPGGVR absolute abundance in HEK293T cells transfected with FMRpolyG_100_. Bars represent mean ± SD. N=2. **C**. Western blot of FMRpolyG from HEK293T cells transfected with the indicated constructs. GAPDH serves as a loading control. **D**. The distribution of two different initiation fragments quantified via PRM-SIS rom IPed lysates of 293Ts transfected with the indicated constructs. **E**. Quantification of the absolute amount of signature peptides PRM-SIS from IPed lysates of 293Ts transfected with the indicated reporters. Detected indicates observed, but not quantified (See supplemental Figure 5)..

As Ac-TEAPLPGGVR is acetylated (a common posttranslational modification of N-terminal amino acids) and the codon 5’ to the ACG is a GUG, coding for valine, we can conclude that this is an N-terminal initiation fragment and not a trypsin cleavage product of a spurious upstream initiation event (Verkerk et al. 1993; Lange et al. 2014; Ree, Varland, and Arnesen 2018). It is possible that Ac-TEAPLPGGVR is derived from the same near cognate codon ACG, which is supported by a similar observation in another gene, *CLK2 (Na et al. 2018)*. However, Ac-TEAPLPGGVR could also arise from Met initiation at the GUG followed by N-terminal Met (iMet) excision, an event that is particularly common when Thr is the second amino acid (Varland, Osberg, and Arnesen 2015; Martinez et al. 2008; Meinnel, Peynot, and Giglione 2005). To determine if Ac-TEAPLPGGVR was due to initiation at the 5’ GUG with Met-tRNA_i_^Met^ followed by iMet excision, we designed two new reporters. In the first, we mutated the GUG upstream of the ACG to GGG. Importantly, this mutation did not alter the strength of the Kozak sequence for the ACG codon. In the second, we mutated the GUG upstream of the ACG to an AUG. We hypothesized that if Ac-TEAPLPGGVR was derived from the ACG, the GGG mutation (mut1) would have no effect on Ac-TEAPLPGGVR abundance, while the AUG mutation (mut2) would eliminate Ac-TEAPLPGGVR. Conversely, if Ac-TEAPLPGGVR was derived from initiation at the GUG upstream of the ACG, the GGG mutation would eliminate Ac-TEAPLPGGVR while the AUG mutation would enhance Ac-TEAPLPGGVR. We observed almost complete loss of Ac-TEAPLPGGVR from our mut1 GGGACG reporter, and a significant increase in Ac-TEAPLPGGVR from our mut2 AUGACG reporter. (Figure 3C-E). We also detected Ac-MTEAPLPGGVR from our mut2 AUGACG (Supplemental Figure 5). These results strongly suggest Ac-TEAPLPGGVR is generated by GUG mediated Met-tRNA_i_^Met^ initiation followed by iMet excision, and identifies a previously overlooked near cognate codon utilized for FmrpolyG synthesis.

### Impact of integrated stress response activation on initiation codon usage for FMRpolyG

During the integrated stress response (ISR), canonical translation initiation factor eIF2α is phosphorylated, preventing new TC formation, and consequently inhibiting global canonical translation (Walter and Ron 2011; Jackson, Hellen, and Pestova 2010). However, CGG RAN translation is selectively enhanced by the ISR in a codon specific manner (Green et al. 2017). As our data supports the role for two different initiation codons, GUG and ACG, we explored whether the ISR was preferentially promoting initiation at one or both of these codons. Similar to what we see with RAN translation-specific nanoluciferase reporters, we observed an ∼2 fold increase in FMRpolyG-3xFlag upon treatment with the ER stress inducer, thapsigargin (TG) (Green, 2017, Figure 4A-B and Supplemental Figure 7A). Using PRM-SIS quantification, both methionine and threonine mediated RAN translational initiation were significantly enhanced by TG (Figure 4C). However, the relative ratio of Ac-TEAPLPGGVR to Ac-MEAPLPGGVR was enhanced by Thapsigargin (Figure 4D). This suggests that GUG mediated initiation in particular may occur independent of the eIF2 ternary complex.

**Figure 4.**
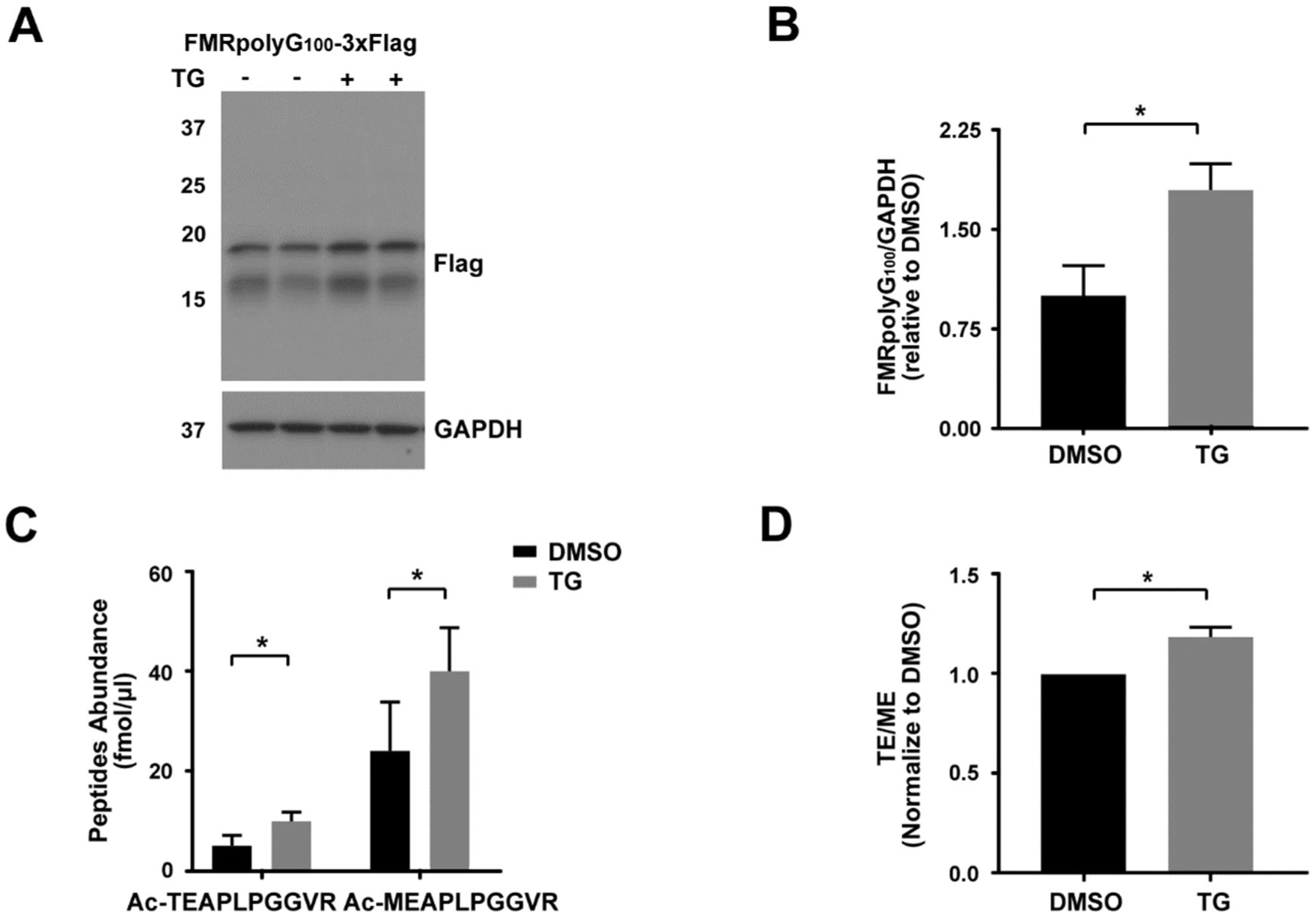
GUG initiation preferentially increases in response to thapsigargin. **A**. Representative western blot of FMRpolyG from HEK293T cells transfected with FMRpolyG_100_-3XFlag followed by DMSO or TG treatment. GAPDH serves as a loading control. N=3. **B**. Quantification FMRpolyG from the above western blot. Bars represent mean ± SD, N=3. *p < 0.05. **C**. The absolute abundance of Ac-TEAPLPGGVR and Ac-MEAPLPGGVR by PRM-SIS in lysates of above samples. Bars represent mean ± SD, N=3. *p < 0.05. **D**. Ratios of the two N-terminal peptides in TG vs DMSO treated 293Ts. Bars represent mean ± SD, N=3. *p < 0.05.

### FMRpolyG synthesis via GUG initiation is partially dependent on eIF2A and inhibited by eIF1 overexpression

Canonical translation initiation is dependent on eIF2 forming a ternary complex with GTP and the initiator Met-tRNA_i_^Met^, which binds the 40S ribosome along with other translation initiation factors to form the PIC. The PIC then binds the 5’ cap of the mRNA and scans along in a 5’ to 3’ direction until it reaches an AUG start codon (Sonenberg and Hinnebusch 2009; Hinnebusch 2011; Jackson, Hellen, and Pestova 2010; Hershey 2000). However, during ISR activation, functional eIF2 is depleted, suggesting that initiation at these near cognate codons likely requires an eIF2 alternative factor. One particular factor, eIF2A, is known to be upregulated during cellular stress (Kim et al. 2018; Ventoso et al. 2006; Starck et al. 2016) and was recently suggested to play a role in RAN translational initiation at G_4_C_2_ repeats in *C9orf72* (Sonobe et al. 2018). Thus we investigated whether eIF2A was required for ACG or GUG initiated RAN translation. We performed PRM-SIS on WT and eIF2A KO HAP1 cell lines transfected with FMRpolyG_100_-3xFlag. We found loss of the alternative eIF2 factor, eIF2A, reduced total FMRpolyG, but significantly suppressed GUG initiation by ∼40%, relative to ACG mediated initiation (Figure 5A-C, Supplemental Figure 6A, and Supplemental Figure 7B). This suggests that GUG initiation is at least partially mediated by eIF2A.

**Figure 5.**
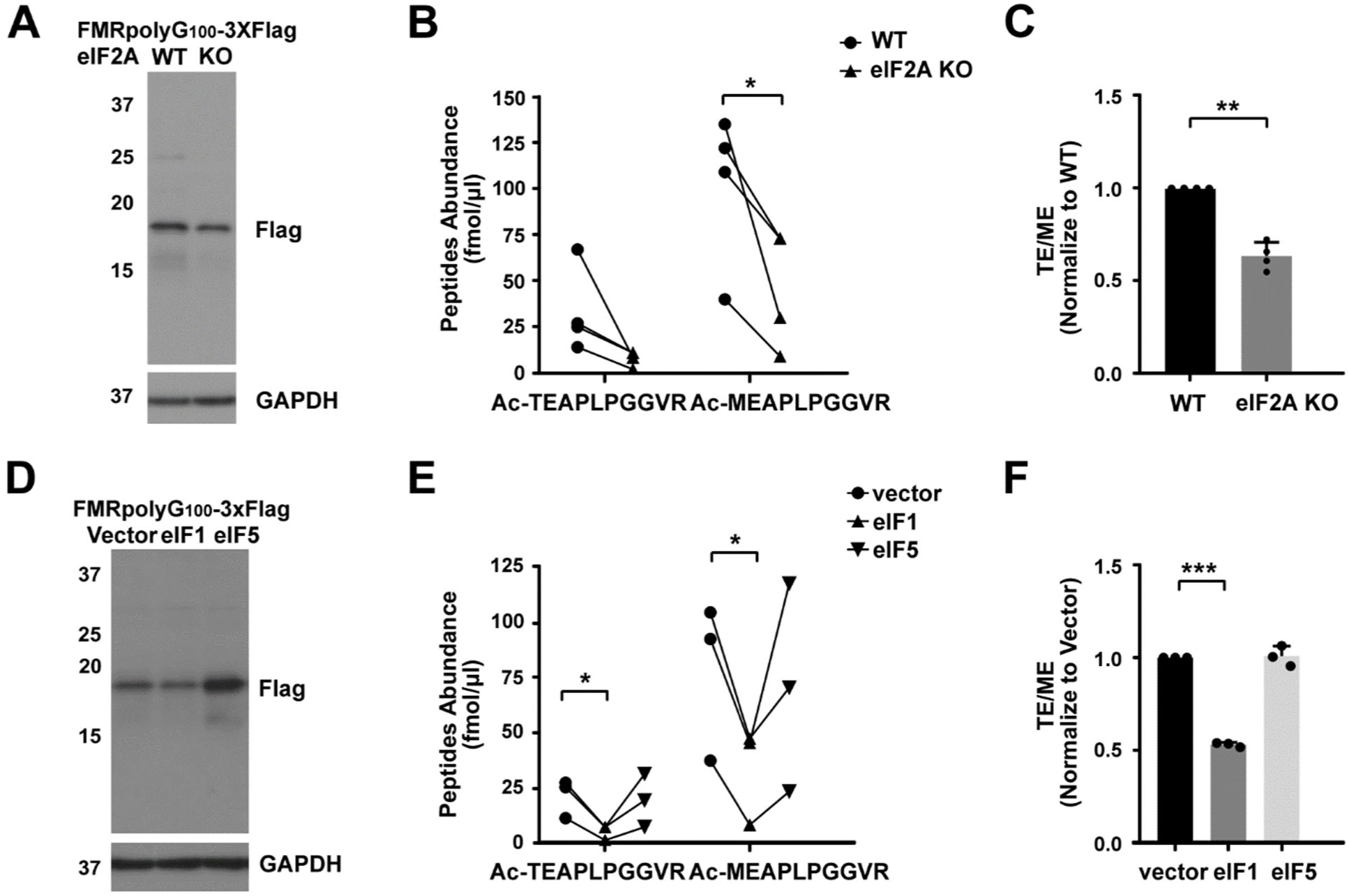
FMRpolyG initiation codon choice is influenced by eIF1 and eIF2A availability. **A**. Representative western blot of FMRpolyG from HAP1 WT and eIF2A KO cells transfected with FMRpolyG_100_-3XFlag. GAPDH serves as a loading control. N=4. **B**. Quantification of Ac-TEAPLPGGVR and Ac-MEAPLPGGVR absolute abundance by PRM-SIS in Flag-IPed lysates of HAP1 cells transfected with FMRpolyG_100_-3XFlag. N=4. *p < 0.05. **C**. Ratio of the two N-terminal peptides in HAP1 WT and eIF2A KO cells transfected with FMRpolyG_100_-3XFlag. Bars represent mean ± SD, N=4. **p < 0.01. **D**. Representative western blot of FMRpolyG from HEK293T cells co-transfected with FMRpolyG_100_-3XFlag and control, eIF1 or eIF5 vectors. GAPDH serves as a loading control N=3. **E**. Quantification of Ac-TEAPLPGGVR and Ac-MEAPLPGGVR absolute abundance by PRM-SIS in Flag-IPed lysates of HEK293T cells co-transfected with FMRpolyG_100_-3XFlag and control, eIF1, or eIF5 vectors. N=3. *p < 0.05. **F**. Ratio of the two N-terminal peptides in HEK293T cells co-transfected with FMRpolyG_100_-3XFlag and control, eIF1 or eIF5 vectors. Bars represent mean ± SD, N=3. ***p < 0.001.

Noncanonical translation initiation can also be facilitated by altered levels of canonical translation initiation factors. In particular, eIF1 and eIF5 have known roles in regulating start codon fidelity, where eIF1 enhances codon stringency, and eIF5 promotes dissociation of eIF1, reducing codon stringency (Pestova and Kolupaeva 2002; Pisarev et al. 2006; Luna et al. 2012). Our lab has previously shown that CGG RAN translation decreases with eIF1 overexpression and increases with eIF5 overexpression (Linsalata et al. 2019). To determine whether eIF1 and/or eIF5 differentially affect ACG or GUG initiation, we co-expressed FMRpolyG_100_-3xFlag with either empty, eIF1, or eIF5 expression vectors, and performed PRM-SIS. While there was no effect on either Ac-MEAPLPGGVR or Ac-TEAPLPGGVR product abundance with eIF5 overexpression, we found overexpression of eIF1 significantly decreased both ACG and GUG initiation, with the impact being greater on GUG initiation, decreasing ∼50% relative to ACG initiation (Figure 5D-F, Supplemental Figure 6B-C and Supplemental Figure 7C). These data are consistent with eIF1 having a preferential effect on GUG initiation.

## Discussion

RAN translated proteins have emerged as important potential participants in the pathogenesis of multiple neurodegenerative disorders. Accurate quantification of these proteins and a deep understanding of the mechanisms underlying their production are both key steps in biomarker and therapy development for these untreatable diseases. Here, using a highly sensitive MS method, PRM-SIS, we identified N-terminal and C-terminal peptides of FMRpolyG in transfected samples and in patient tissue and cells. By comparing endogenous values to their respective SIS, we were able to quantify the absolute amount of FMRpolyG in patient brain tissue, uncovering a large range of FMRpolyG detection in patients. Moreover, this MS based methodology allowed us to identify a novel GUG initiation site for FMRpolyG production and discern its relative usage in conditions of stress and in response to specific perturbations in initiation factor abundance. Together, these data provide new insights into RAN translation and the pathogenesis of FXTAS and other repeat expansion disorders.

Both IRES and ribosomal scanning-dependent initiation mechanisms have been suggested for RAN translation at different repeat expansions (Zu et al. 2011; Zu et al. 2020; Cheng et al. 2018; Sellier et al. 2017; Kearse et al. 2016; Green et al. 2017). In our own work, we previously defined contributions from all near-cognate codons 5’ to the repeat in *FMR1* by mutational analysis, which identified 2 key codons as likely drivers of FMRpolyG production (Kearse et al, 2016). Here we identify a third near-cognate codon as a meaningful contributor to FMRpolyG production. This particular GUG codon was originally dismissed as a contributor because its mutation placed the neighboring ACG codon into a stronger Kozak sequence (Kearse et al. 2016). How initiation site choice is made at these near-cognate codons on *FMR1* is not fully understood, but our data here provide some insight. We observe that loss of eIF2A dramatically reduces RAN translation from both ACG and GUG initiation, with the effect being particularly dramatic for GUG initiation, reducing it 2-fold relative to ACG initiation (Figure 5C). To date, there is no documentation of the effects of eIF2A on near cognate codons ACG and GUG, and thus, the mechanism by which eIF2A makes this distinction is not clear. eIF2A has previously been implicated in Met-tRNA_i_^Met^ mediated initiation at CUG and UUG codons (Starck et al. 2016; Starck et al. 2012; Liang et al. 2014), which share more similarity to GUG than ACG, and could explain the preferential decrease in GUG initiation in our case.

We also observe that GUG initiation is enhanced more so than ACG initiation upon activation of the integrated stress response. There are a number of likely reasons for this. First, eIF2A expression has been shown to increase in response to cellular stress (Starck et al. 2016; Kim et al. 2018; Reid et al. 2014). Given that both GUG and ACG mediated initiation are enhanced during thapsigargin-induced cellular stress, this favors a role for eIF2A in stress-induced CGG RAN initiation. However, both GUG and ACG initiation still occur to some extent in eIF2A knockout cells, suggesting additional alternative initiation factors are at play. Alternatively, this differential stress response may reflect differential recruitment of the TC following 40S ribosomal scanning. ACG has the second lowest free energy of any near cognate codon (after CUG), and thus in general is more efficient at initiating under non-stressed conditions (Lind and Aqvist 2016). However, binding of the TC to a ribosome already sitting on an initiation codon is much stronger and more efficient when the ribosome is on a GUG vs an ACG (Kolitz, Takacs, and Lorsch 2009). Therefore, under non-stressed conditions, when the TC is bound to the PIC, ACG is the more favorable codon; however, under stress, when TC is limited, and recruitment of TC following PIC loading and scanning is necessary, GUG initiation becomes more favorable. Similarly, reduced GUG initiation in the setting of eIF1 overexpression is consistent with previous in vitro work showing eIF1 primarily acts by discriminating codons based on the first position (Lind and Aqvist 2016). In this system, eIF1 levels had no influence on ACG initiation while dramatically altering the free energy state of PIC binding to GUG. eIF1 overexpression has also been shown to strongly repress translation from a GUG but not an ACG codon, particularly when eIF1 expression is not auto regulated, which is the case in our system (Ivanov et al. 2010; Linsalata et al. 2019; Stewart et al. 2015).

A current paradox in the field is the ability to readily detect FMRpolyG by IHC in FXTAS cases and model systems while observing comparably low levels by MS (Ma et al. 2019; Krans et al. 2019; Green et al. 2017; Todd et al. 2013; Kearse et al. 2016). Overall, our PRM-SIS methodology exhibited excellent quantification characteristics in both CSF and cell lysates, with limits of detection for specific FMRpolyG peptides in the sub-nanomolar range. Moreover, immunoprecipitation led to enrichment of FMRpolyG peptides in a linear fashion from transfected cells (Figure 2B and Figure 3B), suggesting that the biochemistry of the peptides is not altered at higher concentrations. This strategy led to reliable detection and quantification of FMRpolyG in both FXTAS patient brains and cells, confirming the presence of FMRpolyG in FXTAS patient tissue and demonstrating definitively that initiation at near-AUG codons above the repeat contributes to its production in people. However, despite its sensitivity, we were not able to measure FMRpolyG using this methodology in CSF and its total abundance in FXTAS patients remained quite low. The exact reasons that this strategy failed to provide a reliable biomarker are not yet clear. Efforts to further increase FMRpolyG enrichment in PRM-SIS MS through enzymatic digests of the polyglycine tract with lysostaphin or 8M urea extraction methods were unsuccessful. It is likely that our immunoprecipitation strategy failed to capture all of the endogenous FMRpolyG protein, given that additional near-AUG codons (i.e. at the GUG closest to the repeat (Figure 1A)) or initiation within the repeat itself can be used for production of isoforms of FMRpolyG whose N-terminus is too short for MS detection and/or lacks the full antibody epitope. To date, C-terminally targeted antibodies to FMRpolyG have not been sufficiently robust to allow for effective immunoprecipitation of this endogenous protein from brain tissue (Todd et al, 2013, Sellier et al, 2017, Krans et al, 2019, and data not shown). The total abundance of FMRpolyG may also be lower than that observed by prior ICC and IHC measurements and reporter assays. Regardless, while our data demonstrate that PRM-MS is capable of reliably quantifying FMRpolyG at high abundance, the low level of detection in endogenous samples indicates that alternative approaches will be needed if this protein is to serve as a viable biomarker in FXTAS.

In sum, the studies presented here describe a new tool for quantifying the non-AUG initiated CGG repeat product, FMRpolyG and provides novel insights into the mechanisms underlying CGG RAN translation, including a specific role for a previously overlooked initiation codon. Furthermore, it describes differential effects on initiation at these codons, highlighting the importance of targeting multiple factors to successfully downregulate RAN translation. Further work into whether these alternative N-terminal ends influence the physical properties of FmrpolyG and its cellular toxicity are warranted, as are efforts to measure RAN translation products as a biomarker of FXTAS and other repeat disorder pathogenesis.

## Methods

### 1. Cell lines and culture

HEK293Ts were obtained from ATCC and grown in high glucose DMEM with 10% fetal bovine serum (FBS), 1% penicillin/streptomycin (P/S). WT and eIF2A KO Hap1 cells were obtained from Horizon Discovery and grown in IMDM with 10%FBS, 1% P/S. All cell lines were maintained in 10 cm dishes at 37°C.

### 2. Reporters

FMRpolyG_100_-3XFlag, and FMRpolyG_100_ RAN translation reporters were made by cloning in their respective sequences into pcDNA3.1+ between BamHI and PspOMI (Supplemental Table 3). eIF1 and eIF5 expression vectors are previously published (Linsalata et al. 2019; Stewart et al. 2015).

### 3. Transfection and treatment

HEK293Ts were seeded at 4.0 × 10^6^ cells/10 cm dish and transfected at ∼60-70% confluency for 24 hrs with 10 µg of either pcDNA3.1+ vector, FMRpolyG_100_ or FMRpolyG_100_-3XFlag and 3:1 Fugene HD to DNA. For stress induced experiments, the cells were transfected as above for 19 hours followed by 5 hrs with specified drugs (Tg, DMSO). For eIF1 and eIF5 experiments, HEK293Ts were seeded as above and cotransfected for 24 hrs with 2.5 µg FMRpolyG_100_-3XFlag and 25 µg of either pcDNA3.1+ vector, eIF1 or eIF5 and 3:1 Fugene HD to DNA.

Hap1 WT and eIF2A KO cells were seeded at 7.0 × 10^6^ cells/10 cm dish and transfected at ∼60-70% confluency for 24 hrs with 10 µg of FMRpolyG_100_-3XFlag and 3:1 Genjet to DNA.

### 4. Clinical Specimens

FXS iPSCs were obtained from Wicell (Doers et al. 2014). Control 2E iPSC line was obtained from the Parent lab at the University of Michigan and have been previously published (Rodriguez et al. 2020). FXTAS peripheral lymphocytes (LCL1) were kindly gifted to us by Stephanie Sherman, and FXTAS skin fibroblasts (F3), and FXS skin fibroblasts (4026) were kindly gifted by Paul Hagerman. All have been previously reported (Garcia-Arocena et al. 2010; Todd et al. 2010). The 3 FXTAS patient frozen brain tissues (FXTAS1-1906, FXTAS2-345, FXTAS3-1558), 2 control frozen brain tissues (control1-382, control2-8662) were obtained from the University of Michigan Brain Bank. FXTAS patient CSF samples were obtained from Rush University Children’s Hospital, and control CSF pool was purchased from Biochemed Services (3057BF). The information for these human derived samples are presented in Supplementary Table 2.

Lymphocyte cells were grow in 1640 RPMI containing 10% FBS and 1% P/S. Skin fibroblasts were grown in high glucose DMEM with 10% FBS, 1% non-essential amino acid (NEAA) and 1% P/S. The CSF and brain tissues were kept in frozen in -80 until use.

### 5. Immunoprecipitation (IP) and silver staining gel

Samples were immunoprecipitated according to manufacturer’s protocol (LSKMAGAG02, Millipore). In brief, cells were washed with 1XPBS 2x then lysed with RIPA buffer (PI89900, Fisher) with Complete™, Mini Protease Inhibitor Cocktail (11836153001, Sigma). Protein concentration for each sample was determined by Pierce™ BCA protein assay kit (23225, Thermo Scientific™). 1.5-4.0 mg of lysate from FMRpolyG_100_ transfected cells or 1.5-5mg of human derived samples were incubated with NTF1 antibody overnight at 4°C followed by adding PureProteome protein A/G mix magnetic beads for another 2 hours. 1.5-4.0 mg of lysate from FMRpolyG_100_-3XFlag transfected cells were incubated with anti-flag M2 magnetic beads (M8823, Sigma) overnight at 4°C. Beads were washed 3x with 1XTBS for 5 minutes at 4°C. IPed proteins were isolated by directly digesting on the beads themselves for mass spectrometry.

### 6. Ultracentrifuge

Samples were centrifuged at 70,000g for 1 hour. The pellet was resuspended in 8M urea, and both the supernatant and the pellet were run on a 15% SDS-PAGE, and blotted with the indicated antibodies.

### 7. Enzymatic digests

For trypsin experiments, lysates from 293Ts transfected with FMRpolyG_100_ or FMRpolyG_100_-3XFlag (dilute to 1 μg/μl) were digested with indicated amounts of trypsin (1 μg/μl) at 37°C for 1 hour). For lysostaphin (LS) experiments, lysates from 293Ts transfected with FMRpolyG_100_ were digested with indicated amounts of LS at room temperature for 1 hour.

### 8. Western blot (WB)

Western blots were performed according to standard protocol (Krans, 2019). The following antibodies were used for western blots where indicated: NTF1 (1:1000, rabbit (Krans et al. 2019)), Flag M2 (1:1000, mouse, F1804, Sigma), eIF2A (1:8000, rabbit, 11233-1-AP, ProteinTech), eIF1 (1:1000, rabbit, 12496S, CST), eIF5 (1:1000, rabbit, 13894S, CST), tubulin-β (mouse, 1:1000, E7, DSHB), and GAPDH (mouse, 1:2000, sc-32233, SCBT) in 5% milk. HRP-conjugated secondary antibodies, goat anti-mouse HRP (111-035-146, Jackson ImmunoResearch), goat anti-rabbit HRP (111-035-144, Jackson Immuno Research) were used at 1:5000 in 5% milk.

### 9. Sample digestion for MS

The silver stained gel slice was destained according to the manufacturer’s protocol (PROTSIL2-1KT, Sigma). The digestion for gel slice, whole lysates and IPed beads were done according to previously published protocols (Singh et al. 2018).

### 10. Liquid Chromatography Mass Spectrometry (LC-MS/MS)

A hybrid quadrupole-orbitrap mass spectrometer (Q Exactive HF, Thermo Scientific) coupled to nano-UHPLC (Ultimate 3000 RSLC Nano, ThermoScientific) was used for acquiring all mass spectrometry data. For Ultra High-Performance Liquid Chromatography (UHPLC), the procedures were according to protocol (Casado et al. 2018).

Resolved peptides were directly sprayed onto the mass spectrometer (MS) operating in either Data-Dependent Acquisition (DDA) or Parallel Reaction Monitoring (PRM, targeted) mode.

For DDA mode, the MS was set to collect one high-resolution MS1 scan (60,000 FWHM @200 m/z; scan range 400-1500 m/z) followed by data-dependent High-energy C-trap Dissociation MS/MS (HCD) spectra on the 20 most abundant ions observed in MS1. Other MS parameters were as previously published (Singh et al. 2018).

For PRM mode: Synthetic peptides with heavy isotope labeled arginine or lysine at the C-termini were obtained from ThermoScientific (AQUA Ultimate grade, 99% pure, ±5% quantitation precision). The digested samples were spiked with 25 fmol/μl of heavy synthetic peptides prior to UHPLC separation. In PRM mode, the quadrupole was set to isolate only the desired precursor ions (see Supplementary Table 1) followed by HCD-MS/MS on the isolated ions.

### 11. Protein identification

Proteome Discoverer (v2.1; Thermo Fisher) was used for data analysis. MS2 spectra were searched against latest SwissProt human protein database (42054 sequences) using the following search parameters: MS1 and MS2 tolerance were set to 10 ppm and 0.1 Da, respectively; carbamidomethylation of cysteines (57.02146 Da) was considered a static modification; acetylation of protein N-terminus, oxidation of methionine (15.9949 Da) and deamidation of asparagine and glutamine (0.98401 Da) were considered as variable modifications. Percolator algorithm (PD2.1) was used to determine the false discovery rate (FDR) and proteins/peptides with ⩽1% FDR were retained for further analysis. MS/MS spectra assigned to the peptides of interest were manually examined.

### 12. Protein quantification by PRM

The raw data was processed using Skyline 4.1.0.18169. A spectral library was created in Skyline for some of the target peptides using several DDA of FMRpolyG samples as well as blanks spiked with AQUA Ultimate endogenous (light) and isotopically labeled (heavy) peptides obtained from ThermoFisher Scientific (Supplementary Table 1). Product ions (minimum of 3) as determined by DDA analysis or Skyline prediction were monitored for each peptide (see the Condensed Transition List.xlsx attachment). Samples analyzed by PRM were spiked with 25 fmol/µl of heavy peptides and the concentration of the endogenous peptide was determined by comparison of the area ratios.

### 13. Statistical analysis

Blots were quantified by Image J software from NIH. Data plots and statistical analyses were done using GraphPad Prism 7. The error bars represent standard deviation (SD) of the mean. Unpaired t-test was used for WB blots intensity. Paired t-test was applied for all PRM-SIS quantification value analysis. We set p < 0.05 as the cutoff for significance.

## Acknowledgements

We thank the University of Michigan Brain Bank, for access to tissues. We thank the University of Michigan Mass Spectrometry-Based Proteomics Resource Facility for their assistance and use of equipment. We thank Amy Krans for cloning plasmids and acquiring brain tissue. We thank Katelyn Green for providing eIF2A KO HAP1 cells, sharing unpublished data and intellectual input.

