## Supplemental Figures and Tables for "Near-cognate initiation generates FMRpolyG from CGG repeats in Fragile X associated Tremor Ataxia Syndrome"

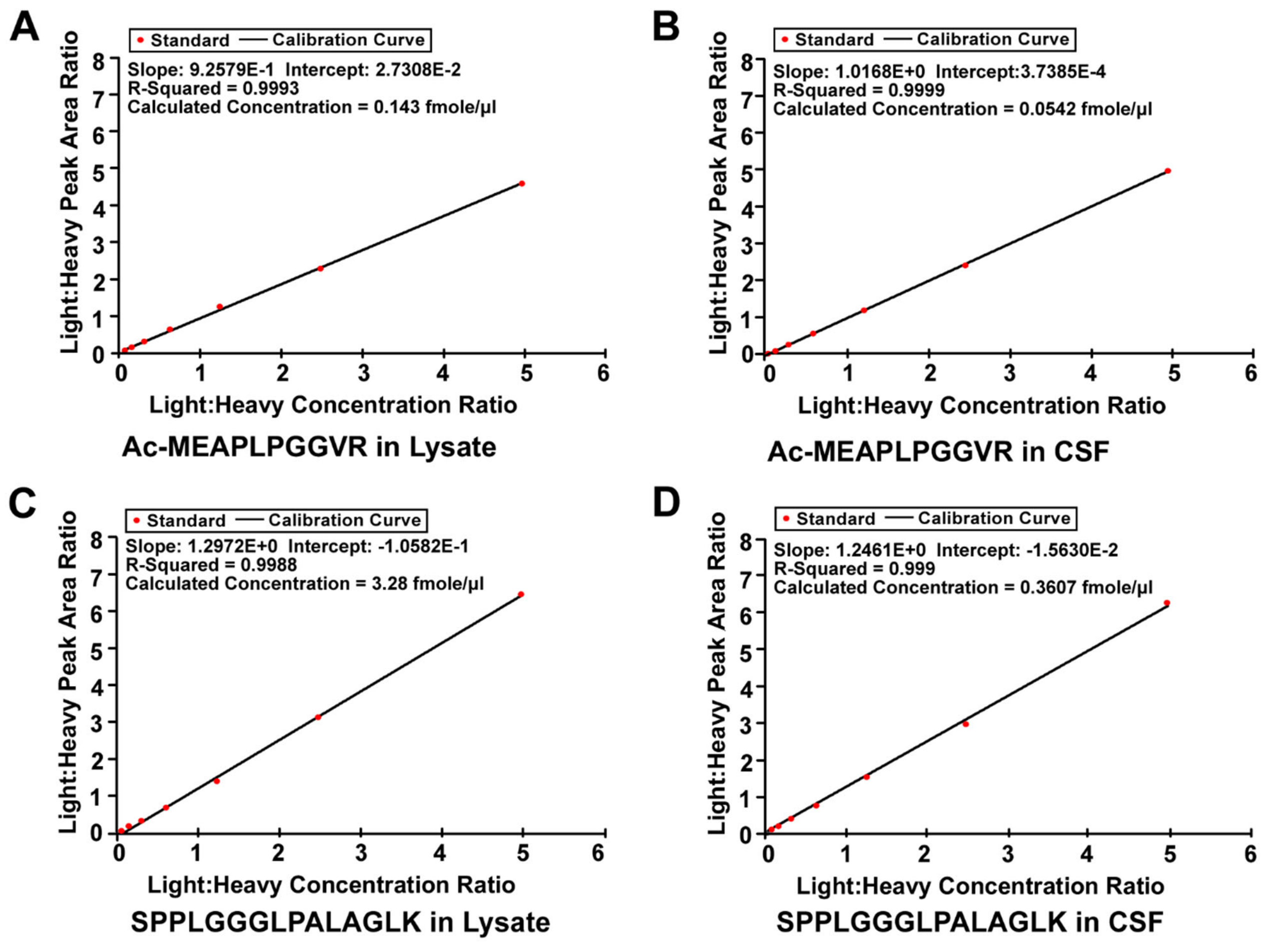


**Supplemental Figure 1. Standard curve for peptides Ac-MEAPLPGGVR and SPPLGGGLPALAGLK of FMRpolyG quantification by PRM-SIS.**

**A, C.** Standard curve for peptides Ac-MEAPLPGGVR (A) and SPPLGGGLPALAGLK (C) in the background of fragile X syndrome patient derived fibroblasts lysate.

**B, D.** Standard curve for peptides Ac-MEAPLPGGVR (B) and SPPLGGGLPALAGLK (D) in the background of control human CSF.

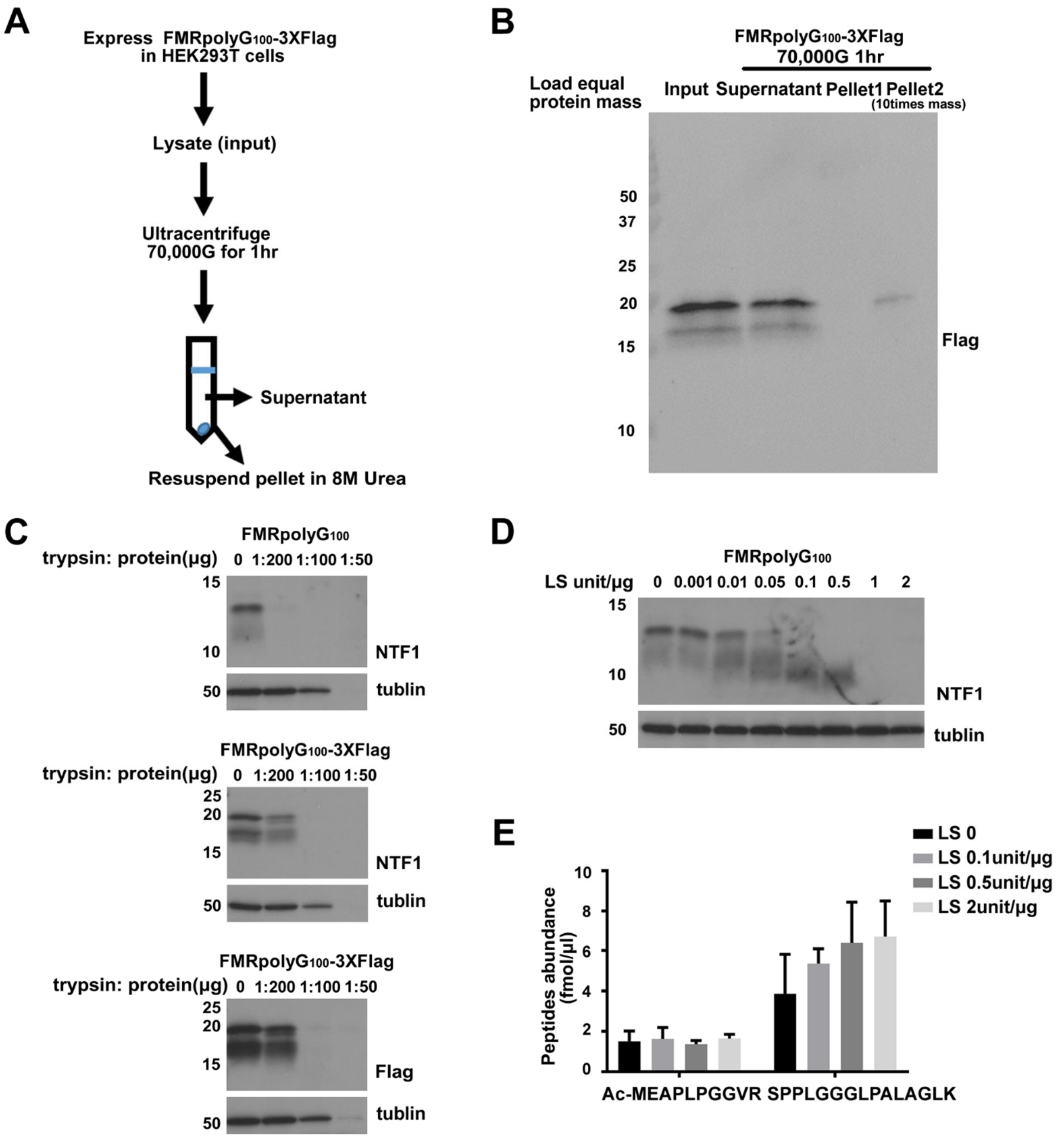


**Supplemental Figure 2. Characterization and optimization of FMRpolyG for PRM-SIS.**

**A-B.** Schematic (A) and western blot (B) of ultracentrifugation and urea treatment for FMRpolyG overexpression in HEK293T lysates.

**C.** Western blot of FMRpolyG from lysates of HEK293T cells transfected with indicated constructs, digested with increasing amounts of trypsin. Top: untagged construct.

**D-E.** Western blots (D) and PRM-SIS quantification (E) of FMRpolyG from lysates of HEK293T cells transfected with untagged FMRpolyG construct after lysostaphin (LS) digestion as indicated concentrations.

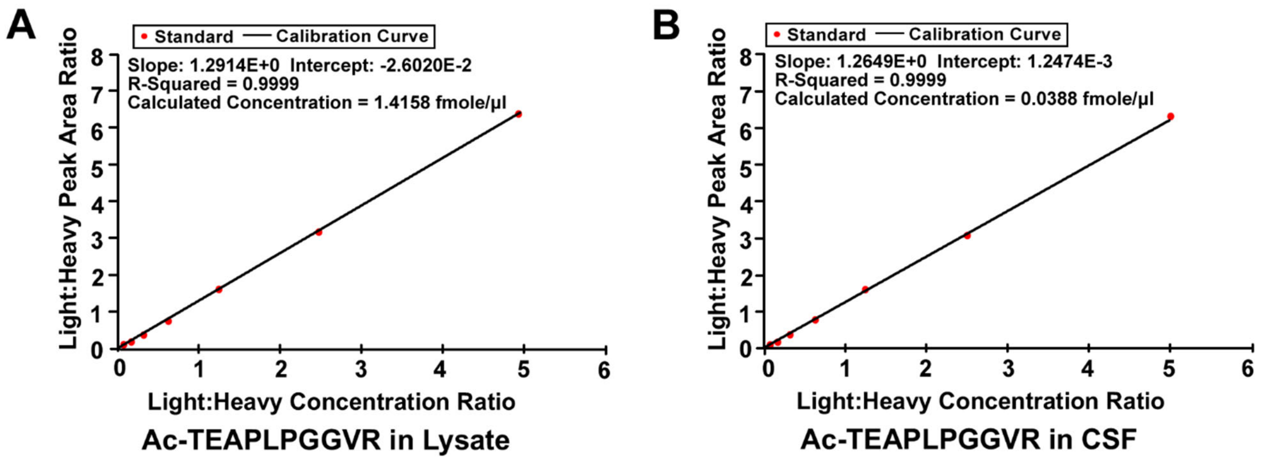


**Supplemental Figure 3. Standard curve for peptide Ac-TEAPLPGGVR of FMRpolyG quantification by PRM-SIS.**

**A.** Standard curve for peptide Ac-TEAPLPGGVR (A) in the background of fragile X syndrome patient derived fibroblasts lysate.

**B.** Standard curve for peptide Ac-TEAPLPGGVR (B) in the background of control human CSF.

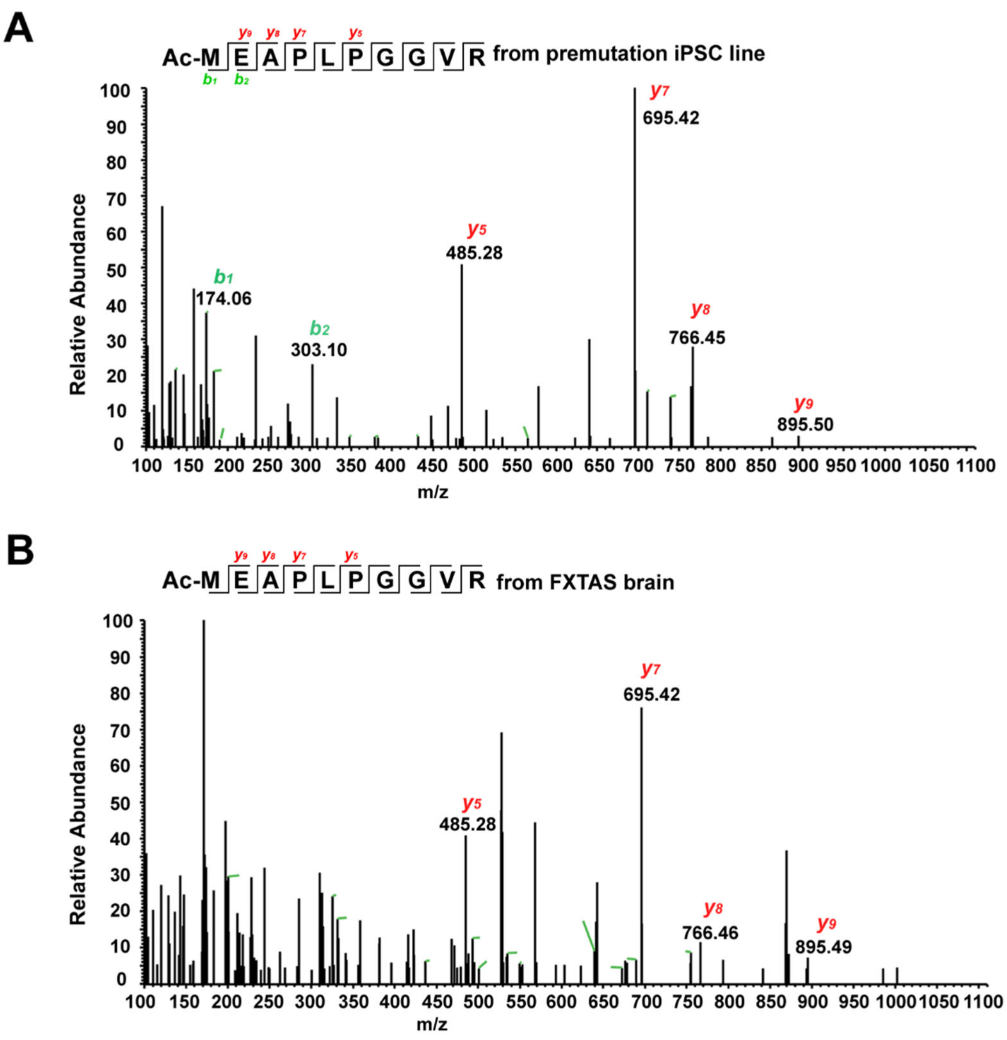


**Supplemental Figure 4: Quantification FMRpolyG from FXTAS patient derived samples by PRM-SIS.**

**A-B.** Representative LC-MS/MS chromatogram of monitored endogenous FMRpolyG peptide Ac-MEAPLPGGVR by PRM-SIS from (A) premutation iPSC line and (B) FXTAS patient brain, following NTF1-IP enrichment. Observed *b-* and *y-ions* are indicated.

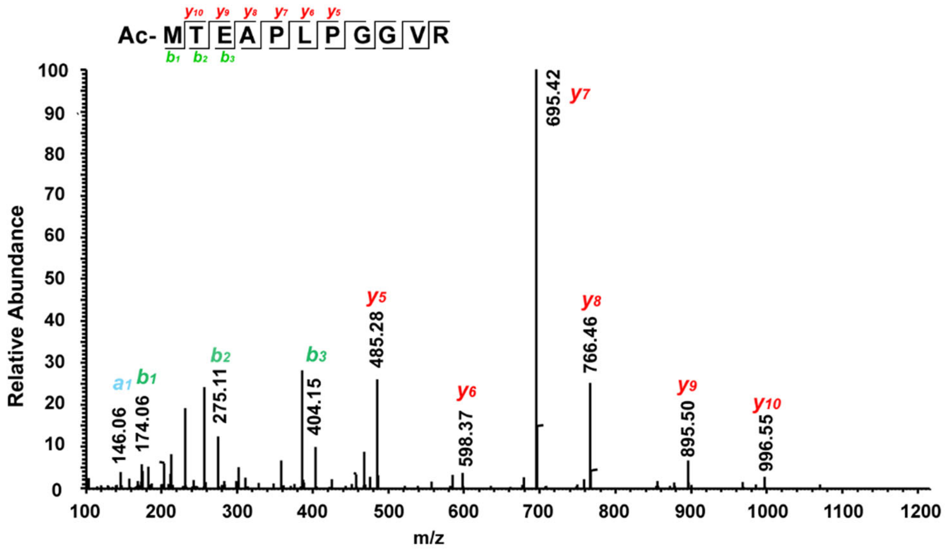


**Supplemental Figure 5: Representative LC-MS/MS chromatogram of monitored Ac-MTEAPLPGGVR peptide.**

LC-MS/MS spectra of Ac-MTEAPLPGGVR from HEK293T cells transfected with mut2 AUGACG reporter, following Flag-IP enrichment. Observed *b-* and *y-ions* are indicated.

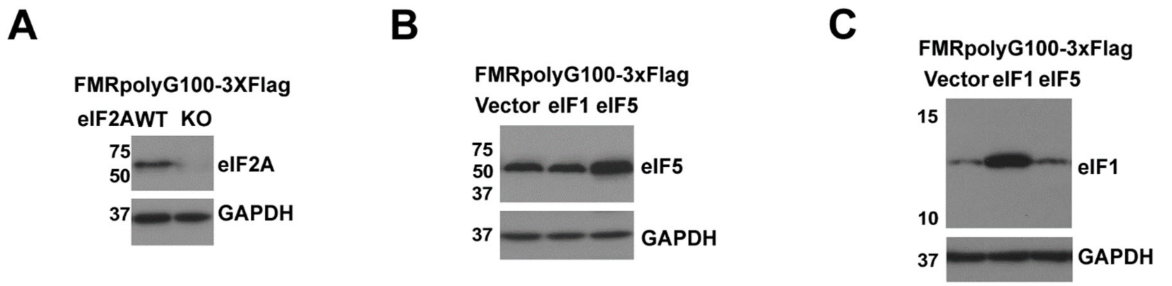


**Supplemental Figure 6. Representative western blots of effects of eIF2A, eIF5 and eIF1 to FMRpolyG expression.**

**A.** Representative western blot of eIF2A from HAP1 WT and eIF2A KO cells transfected with FMRpoly$G_{100}$-3XFlag. GAPDH serves as a loading control.

**B.** Representative western blot of eIF5 from HEK293T cells co-transfected with FMRpoly$G_{100}$-3XFlag and control, eIF1 or eIF5 vectors. GAPDH serves as a loading control.

**C.** Representative western blot of eIF1 from HEK293T cells co-transfected with FMRpoly$G_{100}$-3XFlag and control, eIF1 or eIF5 vectors. GAPDH serves as a loading control.

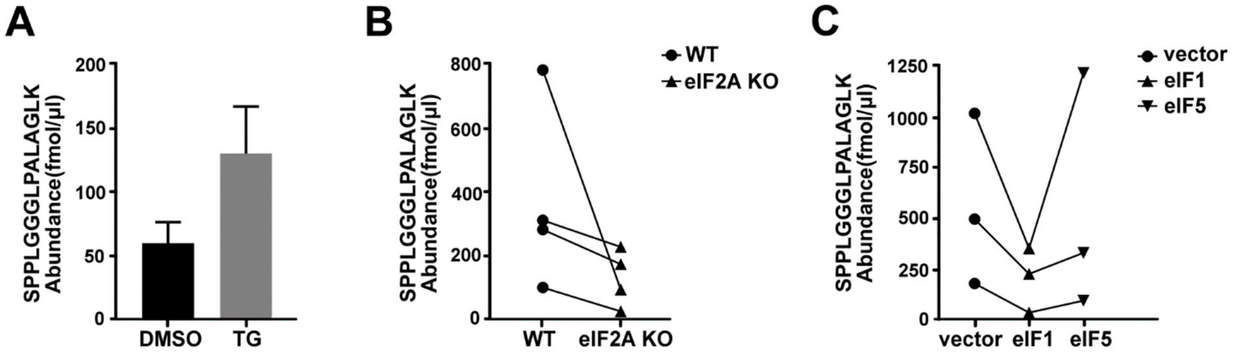


**Supplemental Figure 7. Quantification of SPPLGGGLPALAGLK in different conditions.**

**A.** The absolute abundance of SPPLGGGLPALAGLK by PRM-SIS in lysates from HEK293T cells transfected with FMRpolyG$_{100}$-3XFlag followed by DMSO or TG treatment. Data were shown as mean ± SD, N=3. * indicates $p < 0.05$.

**B.** Quantification of SPPLGGGLPALAGLK absolute abundance by PRM-SIS in Flag-IPed lysates of HAP1 cells transfected with FMRpolyG100-3XFlag. N=4.

Quantification of SPPLGGGLPALAGLK absolute abundance by PRM-SIS in Flag-IPed lysates of **C.** HEK293T cells co-transfected with FMRpolyG100-3XFlag and control, eIF1, or eIF5 vectors. N=3.

**Supplemental Table 1. PRM parameters for monitored peptides for quantification.**

| Monitored peptide | Predicted precursor ion *m/z* with 2 charge | Predicted fragment ion *m/z* for quantification | FXS fibroblasts as background (fmol/µl) | | Human CSF as background (fmol/µl) | |
|---|---|---|---|---|---|---|
| | | | LOD | LOQ | LOD | LOQ |
| Ac-MEAPLPGGVR | 534.779 | 766.457(y8)<br>695.4199(y7)<br>485.2831(y5) | - | | | - |
| SPPLGGGLPALAGLK | 674.4034 | 1163.7147(y13)<br>1066.6619(y12)<br>953.5778(y11)<br>669.4294(y7) | - | | | - |
| Ac-TEAPLPGGVR | 519.7826 | 766.457(y8)<br>695.4199(y7)<br>485.2831(y5) | - | | | - |
| Ac-MEAPLPGGV**(R)** * | 544.779 | 776.457(y8)<br>705.4199(y7)<br>495.2831(y5) | 0.9766 | 2.9298 | 0.0814 | 0.2441 |
| SPPLGGGLPALAGL**(K)***| 682.4034 | 1171.7147(y13)<br>1074.6619()y12)<br>961.5778(y11)<br>677.4294(y7) | 1.9531 | 5.8693 | 0.0814 | 0.2441 |
| Ac-TEAPLPGGV**(R)** * | 529.7826 | 776.457(y8)<br>705.4199(y7)<br>495.2831(y5) | 0.9766 | 2.9298 | 0.0814 | 0.2441 |

*: (R) and (K) are stable isotope labeled, making comprising peptide masses 10Da and 8Da larger than unlabeled equivalents. The limit of detection (LOD) and limit of quantification (LOQ) were determined in two different background, including Fragile X Syndrome patient derived skin fibroblasts and control human CSF.
All the *m/z* information are sourced from Protein Prospector by UCSF from website of http://prospector.ucsf.edu/prospector/mshome.htm.

**Supplemental Table 2. Characteristics for all human derived samples.**

| Samples | Diagnosis | Age(y) | Gender | CGG repeats | Source | Samples' amount used (total protein/ volume) | Quantification of Ac-MEAPLPGGVR by PRM-SIS |
|---|---|---|---|---|---|---|---|
| FX 11-9U | FXTAS | 7 | M | 114 | iPSC line | 1.5mg | 4.23 fmol |
| 2E | Control | - | - | 23 | iPSC line | 1.5mg | ND |
| TC43 | FXS | - | - | un-methylated full mutation | NPC | 7.5mg | 18.0 fmol |
| FXTAS-1 | FXTAS | 74 | M | 102 | Cortex | 5.0mg | 151.2 fmol |
| FXTAS-2 | FXTAS | 78 | M | 90 | Cortex | 3.0mg | 13.41 fmol |
| FXTAS-3 | FXTAS | 80 | M | 90 | Cortex | 3.0mg | 136.08 fmol |
| | | | | | Hippocampus | 4.1mg | 12.15 fmol |
| Control-1 | Control | 81 | M | - | Cortex | 5.2mg | ND |
| Control-2 | Control | 72 | M | - | Cortex | 5.4mg | ND |
| Lym-1 | FXTAS | 60 | M | 114 | Lymphocytes | 3.5mg | ND |
| Fibro-1 | FXTAS | 60 | M | 97 | Fibroblasts | 3.5mg | ND |
| CSF1 | FXTAS | - | - | - | CSF | 5ml | ND |
| CSF2 | FXTAS | - | - | - | CSF | 5ml | ND |
| CSF3 | Control | - | - | - | CSF | 5ml | ND |

ND：not detectable.

**Supplementary Table 3. Vector Sequences**

| Plasmid | Insert Sequence |
|---|---|
| FMRpolyG$_{100}$ | GGATCCACTAGTCCAGTGTGGTGGAATTCGTTAACAGATCTGCTCAGCTCCGTTTCGG<br>TTTCACTTCCGGTGGAGGGCCGCCTCTGAGCGGGCGGCGGGCCGACGGCGAGCGC<br>GGGCGGCGGCGGTGACGGAGGCGCCGCTGCCAGGGGGCGTGCGGCAGCGCGGCG<br>GCGGCGGCGGCGGCGGCGGAGGCGGCGGCGGCGGCGGCGGCGGCGGCGGCGGC<br>GGCGGCGGCGGCGGCGGCGGCGGCGGCGGCGGCGGCGGCGGCGGCGGCGGCGG<br>CGGCGGCGGCGGCGGCGGCGGCGGCGGCGGCGGCGGCGGCGGCGGCGGCGGCG<br>GCGGCGGCGGCGGCGGCGGCGGCGGCGGCGGCGGCGGCGGCGGCGGCGGCGGC<br>GGCGGCGGCGGCGGCGGCGGCGGCGGCGGCGGCGGCGGCGGCGGCGGCGGCGG<br>CGGCGGCGGCGGCGGCGGCGGCGGCGGCGGCGGCTGGCCTCGAGCGCCCGCAG<br>CCCACCTCTCGGGGGCGGGCTCCCGGCGCTAGCAGGGCTGAAGAGAAGATGGAGG<br>AGCTGGTGGTGGAAGTGCGGGGCTCCAATGGCGCTTTCTACAAGGCATTTGAAAGCG<br>GCCGCATAATAGTGAAGTAGGGGCCC |
| FMRpolyG$_{100}$-3xF | GGATCCACTAGTCCAGTGTGGTGGAATTCGTTAACAGATCTGCTCAGCTCCGTTTCGG<br>TTTCACTTCCGGTGGAGGGCCGCCTCTGAGCGGGCGGCGGGCCGACGGCGAGCGC<br>GGGCGGCGGCGGTGACGGAGGCGCCGCTGCCAGGGGGCGTGCGGCAGCGCGGCG<br>GCGGCGGCGGCGGCGGCGGAGGCGGCGGCGGCGGCGGCGGCGGCGGCGGCGGC<br>GGCGGCGGCGGCGGCGGCGGCGGCGGCGGCGGCGGCGGCGGCGGCGGCGGCGG<br>CGGCGGCGGCGGCGGCGGCGGCGGCGGCGGCGGCGGCGGCGGCGGCGGCGGCG<br>GCGGCGGCGGCGGCGGCGGCGGCGGCGGCGGCGGCGGCGGCGGCGGCGGCGGC<br>GGCGGCGGCGGCGGCGGCGGCGGCGGCGGCGGCGGCGGCGGCGGCGGCGGCGG<br>CGGCGGCGGCGGCGGCGGCGGCGGCGGCGGCGGCTGGCCTCGAGCGCCCGCAG<br>CCCACCTCTCGGGGGCGGGCTCCCGGCGCTAGCAGGGCTGAAGAGAAGATGGAGG<br>AGCTGGTGGTGGAAGTGCGGGGCTCCAATGGCGCTTTCTACAAGGCATTTGAAAGCG<br>GCCGCAGACTACAAAGACCATGACGGTGATTATAAAGATCATGACATCGACTACAAGG<br>ATGACGATGACAAGTAAGGGCCC |
